# Genome-wide Mapping of Topoisomerase Binding Sites Suggests Topoisomerase 3α (TOP3A) as a Reader of Transcription-Replication Conflicts (TRC)

**DOI:** 10.1101/2024.06.17.599352

**Authors:** Hongliang Zhang, Yilun Sun, Sourav Saha, Liton Kumar Saha, Lorinc S. Pongor, Anjali Dhall, Yves Pommier

**Affiliations:** Laboratory of Molecular Pharmacology and Developmental Therapeutics Branch, Center for Cancer Research, National Cancer Institute, National Institutes of Health, Bethesda, Maryland, USA

## Abstract

Both transcription and replication can take place simultaneously on the same DNA template, potentially leading to transcription-replication conflicts (TRCs) and topological problems. Here we asked which topoisomerase(s) is/are the best candidate(s) for sensing TRC. Genome-wide topoisomerase binding sites were mapped in parallel for all the nuclear topoisomerases (TOP1, TOP2A, TOP2B, TOP3A and TOP3B). To increase the signal to noise ratio (SNR), we used ectopic expression of those topoisomerases in H293 cells followed by a modified CUT&Tag method. Although each topoisomerase showed distinct binding patterns, all topoisomerase binding signals positively correlated with gene transcription. TOP3A binding signals were suppressed by DNA replication inhibition. This was also observed but to a lesser extent for TOP2A and TOP2B. Hence, we propose the involvement of TOP3A in sensing both head-on TRCs (HO-TRCs) and codirectional TRCs (CD-TRCs). In which case, the TOP3A signals appear concentrated within the promoters and first 20 kb regions of the 5’ -end of genes, suggesting the prevalence of TRCs and the recruitment of TOP3A in the 5’-regions of transcribed and replicated genes.

**GRAPHICAL ABSTRACT:** 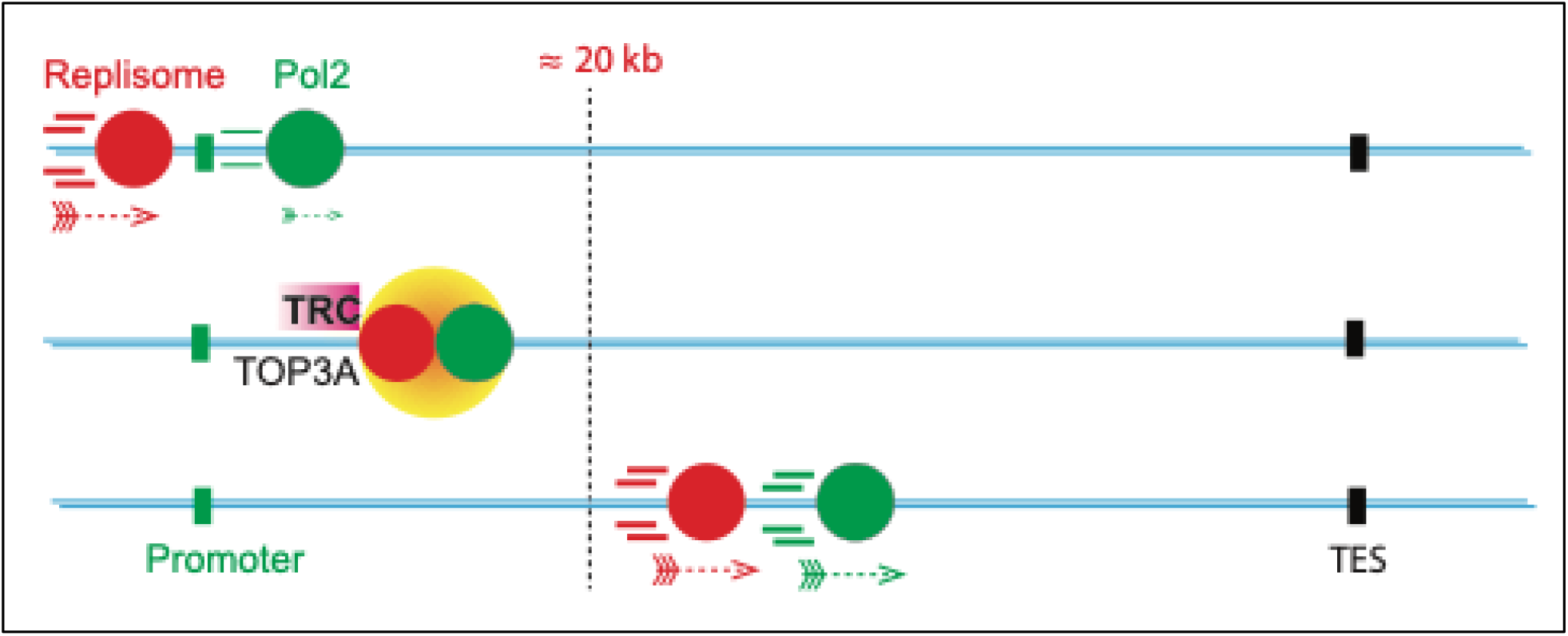

## INTRODUCTION

Topoisomerases play a pivotal role in fundamental nucleic acid metabolic processes^1,2^, including DNA replication, transcription, recombinations and chromatin remodeling. They adjust DNA topology by transiently cleaving and rejoining DNA strands during their catalytic cycle. Depending on whether they transiently cleave one or two strands of DNA they are classified as type I and type II topoisomerases, respectively^3^. The type I topoisomerases are further subcategorized as type IA or IB. Type IA topoisomerases^4^, including human topoisomerase IIIα (TOP3A) and IIIβ (TOP3B) use a strand passage mechanism to carry out their reactions^5^. They cleave single-stranded (unannealed) segments of DNA by generating a covalent linkage to the 5’-phosphate of the DNA backbone in Mg^2+^-dependent and ATP-independent manner, allowing the passage of an intact strand through the cleaved strand. By contrast, the type IB topoisomerases,^6^ including topoisomerase I (TOP1) and the mitochondria-targeted topoisomerase I (TOP1MT), use a controlled rotation mechanism,^7^ whereby they cleave one strand of the duplex DNA backbone by covalently linking the terminal 3’-phosphate, allowing the rotation of the intact strand around the DNA axis and the relaxion of both positive and negative supercoils independently of ATP and Mg^2+^. The type II topoisomerases^5^ are subcategorized as type IIA and IIB. Type IIA topoisomerases including human topoisomerase IIα (TOP2A) and IIβ (TOP2B) produce double-stranded DNA (dsDNA) breaks with canonical 4-base overhangs, while type IIB generate 2-base overhangs. Both type IIA and IIB topoisomerases catalyze dsDNA breaks by cutting the DNA backbone while generating a covalent linkage to the terminal 5’-phosphates on both strands of the DNA duplex in ATP- and Mg^2+^-dependent manner. They change the topology of DNA by a strand passage mechanism at duplex DNA crossovers.

The five topoisomerases (TOP1, TOP2A, TOP2B, TOP3A, and TOP3B) are simultaneously present in the nucleus of a typical human cell to couple the genome topology with the metabolic and structural requirements of the cell^8^. While all topoisomerases change the topology of DNA, the structural and mechanistic differences among them bestow DNA sequence and structure, and activity preferences^9,10^. How topoisomerase activities are distributed across the genome has begun to be addressed using next-generation sequencing (NGS) technologies^11^. Most methods are based on Chromatin ImmunoPrecipitation followed by sequencing (ChIP-seq) to determine genome-wide topoisomerase binding sites or topoisomerase activity as cleavage complexes (TOPccs). Mapping the binding and cleavage sites of TOP1^12^, TOP2A^13^ and TOP2B^14,15^ in mouse and human cells is critical for understanding the biology and functions of topoisomerases and the sites of actions of the widely used anti-cancer drugs targeting TOP1 and TOP2^16–18^. Yet, the signal to noise ratio (SNR) in topoisomerase ChIP-seq data is generally low, making it difficult to fully elucidate the genomic locations of topoisomerase binding and activity. Given the limitations of NGS technologies for topoisomerase genomic mapping, improving signal to noise ratio while maintaining simplicity and consistency of protocols is highly desirable. The present study addresses this point by describing and implementing a modified CUT&Tag method^19^ in cells overexpressing each of the five nuclear topoisomerases. Based on this new approach, we report here the mapping and comparison of the genomic binding sites of the five nuclear topoisomerases (TOP1, TOP2A, TOP2B, TOP3A, TOP3B) and relate their binding to transcription.

Transcription and replication use the same genomic templates and can take place simultaneously leading to transcription-replication conflicts (TRC)^20^. Thus, it is plausible that topoisomerases are implicated in limiting and resolving TRC as converging transcription and replication complexes generate positive supercoiling in front of them and negative supercoiling behind^12,17,21,22^. Depending on the relative directions of transcription and replication, TRC may be head-on (HO) or co-directional (CD)^23^. HO-TRC are the most damaging as the transcription and replication machinery must pass across each other^24,25^. In general, CD-TRC are considered less detrimental than HO-TRC, consistent with the fact that for most genes, especially for the essential and highly transcribed genes, transcription and replication are co-directional^26^. Yet, CD-TRC are still likely to occur when replication is faster than transcription. In bacteria, this could be explained by the fact that replication is 12–30 times faster than transcriptions.^23^ In human cells, the average replication speed is 4.4 kb/min *in vitro*^27^ and 1-2 kb/min^28^ *in vivo*, which can be faster than the transcription speed (0.5-3 kb/min *in vivo*)^29^. Although it is still not established whether the replication machinery can bypass transcription complexes at CD-TRC, evidence suggests that in human cells both CD-TRC and HO-TRC are detrimental to the genome^20,30^. To reduce TRCs and minimize genomic damage, transcription has been shown to take place in isolated membrane-less nuclear bodies^31^, such as the nuclear speckles^32^.

In this study we mapped the genomic sites of potential DNA topological stress by mapping and comparing the chromatin binding of each of the five nuclear topoisomerases in base line conditions and upon replication arrest by the specific replicative polymerase inhibitor, aphidicolin.^33^

## MATERIALS AND METHODS

### Plasmids and site directed mutagenesis

N-terminally FLAG-6×His-tagged TOP1 complementary DNA (cDNA) was amplified by polymerase chain reaction (PCR) using pGALhTOP1 yeast plasmid as a template. The PCR product was inserted into Psp XI/Not I sites of a pT-REx-DEST Gateway vector (Invitrogen). For human TOP2A expression in HEK293 cells, N-terminally FLAG-tagged TOP2A cDNA was amplified by PCR using pMJ1hTOP2A yeast plasmid as a template and then inserted into Psp XI/Not I sites of the pT-REx-DEST Gateway vector. N-terminally FLAG-tagged TOP2B cDNA (derived from pHT500hTOP2B yeast plasmid) was PCR-amplified and inserted into Psp XI/Not I sites of the pT-REx-DEST Gateway vector ^34^. Primers used for the generation of TOP1 and TOP2 expression plasmids are listed in table 1.

**Table 1:**
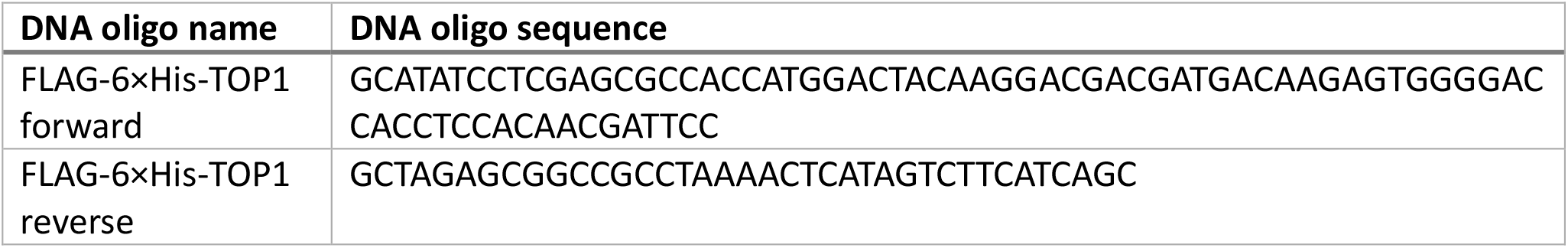
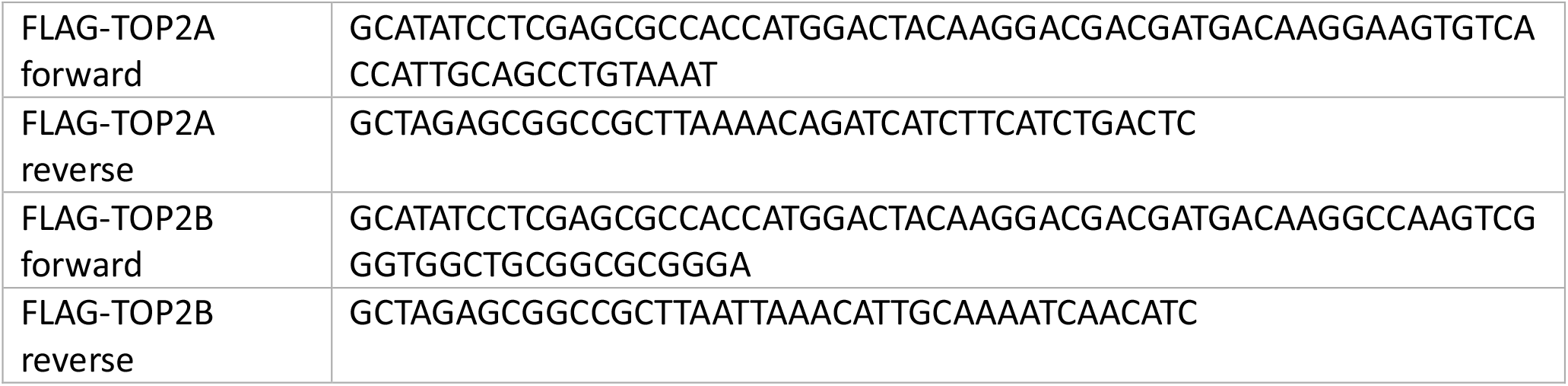
Primers used for the generation of TOP1 and TOP2 expression plasmids.

### Site-directed mutagenesis

The TOP1 T718A/N722H, TOP2A D48N, and TOP2B D64N self-trapping mutants were generated by the Q5 SDM Kit (NEB, catalog no. E0554S) following manufacturer’s instruction and mutation confirmation by sequencing. Primers used for generating the TOP1 and TOP2A and B mutant expression constructs are listed in table 2.

**Table 2:**
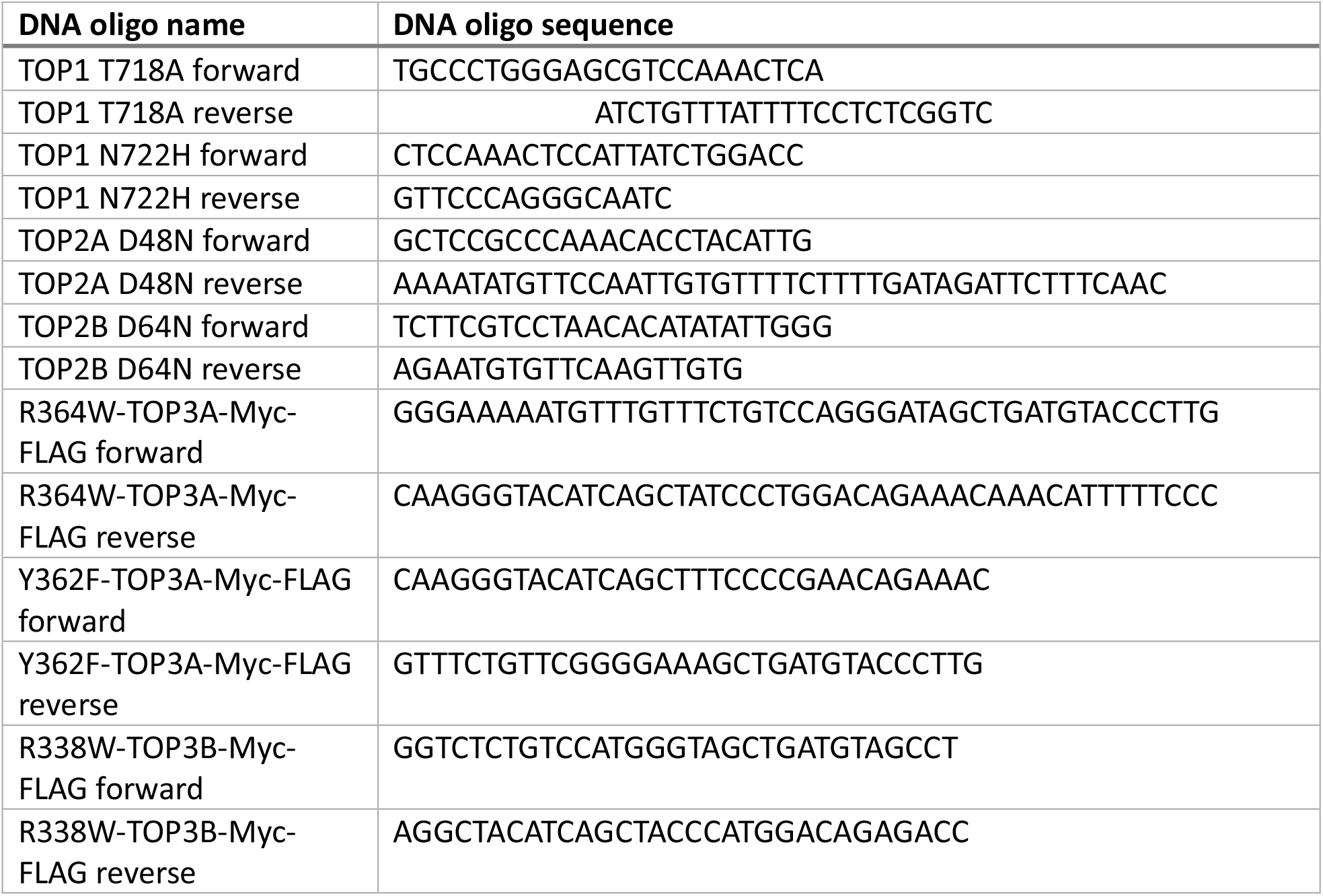
Primers used for generating the topoisomerase mutant expression constructs.

Human TOP3A-Myc-FLAG cDNA ORF (CAT#: RC208236) and human TOP3B-Myc-FLAG cDNA ORF (CAT#: RC223204) Clones were purchased from OriGene. Site-directed mutagenesis was performed using QuikChange II XL site-directed mutagenesis kit (Agilent Technologies) following the manufacturer’s protocol, and mutations were confirmed by sequencing. Primers used for generating the TOP3A^35^ and TOP3B^22^ mutant expression constructs are listed in table 2.

### Cell lines, culture conditions and transfection of expression plasmids

Human HEK293 cells were cultured in Dulbecco’s Modified Eagle Medium (DMEM) (Cat# 084564, Gibco, US) supplemented with fetal bovine serum (10%, Gibco, US), penicillin (100 U/ml), and streptomycin (100 μg/ml, ThermoFischer, US), and maintained at 37 °C under a humidified atmosphere and CO_2_ (5%). Transient transfection of expression plasmids was carried out using Lipofectamine 3000 reagents (CAT#: L3000015, ThermoFischer, US) according to the manufacturer’s protocol for 48 h.

### CUT&Tag method

The pA-Tn5 transposome was prepared following the protocol of Kaya-Okur et al.^36^ The CUT&Tag protocol was carried out following Carter et al^37^ with modification. Briefly, 2 μl 2XCB (0.1 M Tris pH 8.0; 0.3 M NaCl; 0.1% Triton X-100; 25% Glycerol), 2 μl pA-Tn5 tranposome (at 4 μM), and 2 μl antibody (at 0.2 mg/ml for monoclonal antibody, at 0.5 mg/ml for polyclonal antibody) were mixed in a microcentrifuge tube (mixture A) and kept on ice. One million cells were harvested and transferred to a clean 1.5 mL tube. Cells were washed once with 1 ml of PBS, suspend in 1 mL of 1X CB, and incubated on ice for 10 min. The cells were spun down, suspend in 1 mL of 1X CB. An aliquot of 50 μl was added to mixture A, incubated at room temperature for 60 min with rotation. 500 μl of wash buffer (50 mM Tris pH 8.0; 150 mM NaCl; 0.05% Triton X-100) was added to each tube, centrifuged. The pellets were washed two more time with 500 μl of wash buffer, and then suspended in 100 μl of wash buffer. 1 μl of 1 M MgCl_2_ was added to start the transposome reaction, incubated at 37°C for 60 min. 4 μl of 0.5 M EDTA, 2 μl of 10% SDS and 1 μl of 20 mg/mL Proteinase K were added, and incubated at 55 °C for 60 min. The DNA was purified with ChIP DNA clean & concentrator (Zymo Research). Library was constructed following manufacture’s protocol (Illumina) and sequenced on Next-seq.

### Data analysis

The generated fastq files were quality controlled with FastQC (https://www.bioinformatics.babraham.ac.uk/projects/fastqc/), aligned to hg19, hg38, and T2T reference genome with bwa-mem2 (https://github.com/bwa-mem2/bwa-mem2/tree/master), deduplicated, sorted and indexed using Samtools^38^ and Picard (http://broadinstitute.github.io/picard). BigWig files were generated with BAMscale^39^ and visualized using Integrative Genomics Viewer (IGV)^40^. Reads per million (RPM) values were calculated around protein coding genes. The profiles of short reads’ average distribution along normalized gene bodies were generated by ngs.plot^41^. The profiles were smoothed using the sliding window algorithm. The transcription vs topoisomerase signal plots, the control vs aphidicolin-treatment plots were generated with ggplot2^42^ package in R. Correlation heatmap was generated with UCSC genome browser’s “bigWigCorrelate” tool.ggplot2 R package.

Broad peaks for TOP1, TOP2A, TOP2B, TOP3A, and TOP3B were called with MACS2 software (version 2.2.7.1)^43^, and these peaks were used for the downstream analysis. The annotation of peaks in several genomic regions, such as TSS (transcription start site), TTS (transcription termination site), promoter, Exon (Coding), 5’ UTR, 3’ UTR, Intronic, or Intergenic, was generated with HOMER (version 4.11.1) software (http://homer.ucsd.edu/homer). Enrichment analysis performed using Enrichr (pmid: 27141961), it applies Fisher exact test to identify enrichment scores. For the TOP3A log_2_ratio, the formula was log_2_((untreated signal + 1)/(aphidicolin-treated signal + 1)).

## RESULTS

### CUT&Tag method for mapping topoisomerase binding sites and comparison with prior methods

To improve the noise to signal ratio (NSR), we applied two factors into our sequencing: a modification without fixation of the CUT&Tag method (Fig. 1) and transfection of HEK293 cells with FLAG-tagged topoisomerase constructs. We designed the topoisomerase expression constructs based on our recent cellular studies with three different topoisomerase 3α (TOP3A) constructs^35^ and compared the CUT&Tag signals obtained wild-type TOP3A (WT), its active site mutant Y362F containing a tyrosine to phenylalanine replacement at position 362 (YF) and the self-trapping R364W TOP3A mutant^35^ containing an arginine to tryptophan mutation (RW) at position 364. CUT&Tag with the WT, YF and RW forms of TOP3A produced comparable binding patterns including for the YF mutant (Fig. S1A), indicating that the detected binding was mainly non-covalent. Closer analysis showed that TOP3A-R364W generated stronger reads than the wild-type (WT) and the catalytic-dead mutant (YF) (Fig. S1B). So, we chose to generate and map the binding sites of the self-trapping forms for all topoisomerases in our following genome-wide mapping experiments. For TOP1 we generated and used the TOP1-T718A/N722H double-mutant based on sequence homology with our previously reported self-trapping TOP1MT^44^. For TOP2α we constructed the TOP2A-D48N mutant based on a prior study^45^. For TOP2β we constructed the TOP2B-D64N mutant by sequence homology with the TOP2A-D48N mutant. And for the type IA enzymes we used the TOP3A-R364W and TOP3B-R338W self-trapping mutants based on our recent publications^22,35^. Clear genomic signals were obtained for the 5 nuclear topoisomerases (Fig. 2 and Fig. 3).

**Fig. 1.**
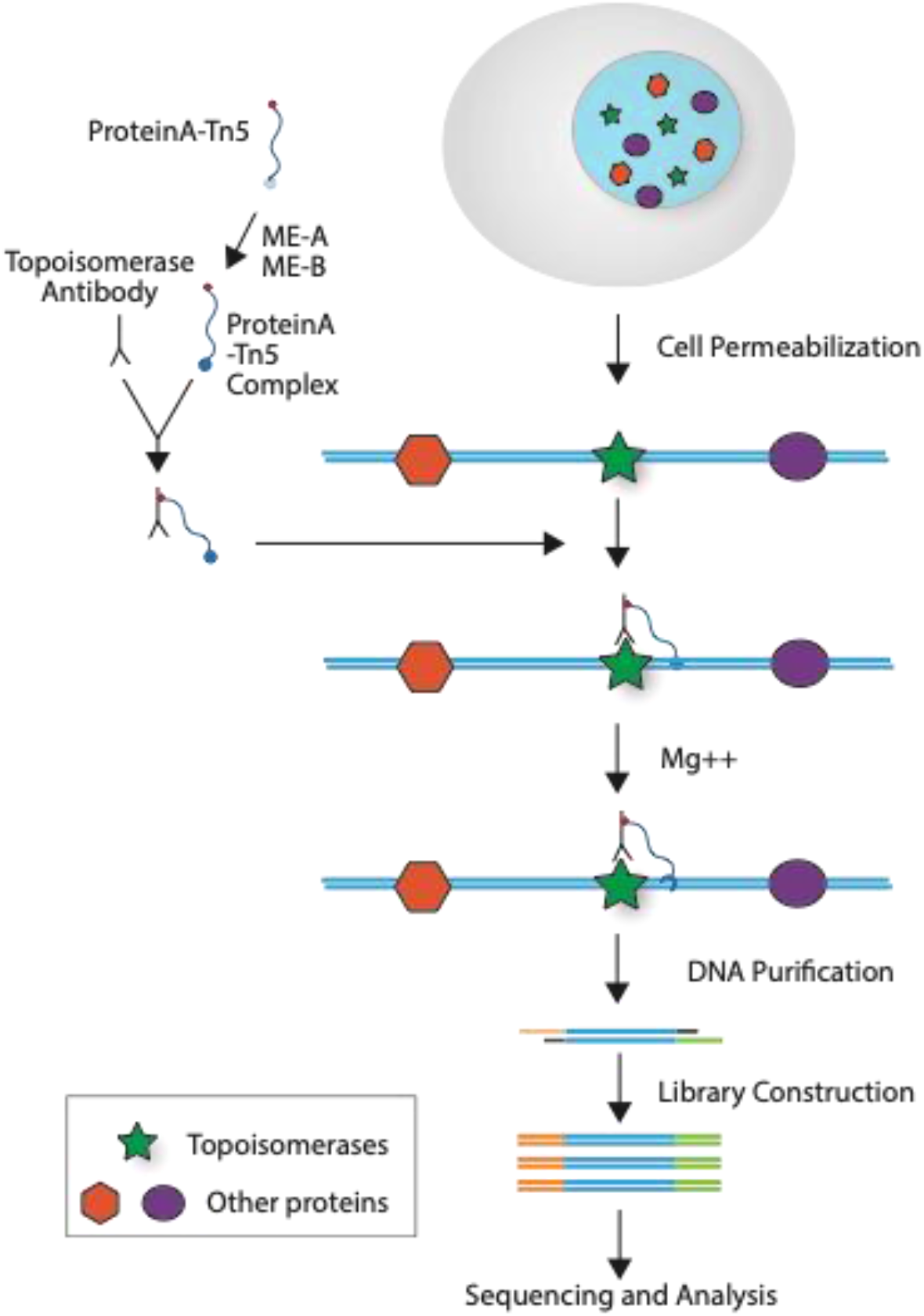
Topoisomerase binding CUT&Tag sequencing: Purified proteinA-Tn5 was incubated with ME-A and ME-B to make the proteinA-Tn5 complex, which was mixed with topoisomerase-specific antibody on ice for 30 minutes. The mixture was incubated with permeabilized cells for 60 minutes at room temperature to allow topoisomerase binding to its specific antibody. Free antibody was washed away and Mg++ was added to start the CUT& Tag process for 60 minutes at 37oC. Reactions were stopped by adding EDTA, SDS, and proteinase K. The DNA was purified and libraries were constructed, followed by sequencing and data analysis.

**Fig. 2.**
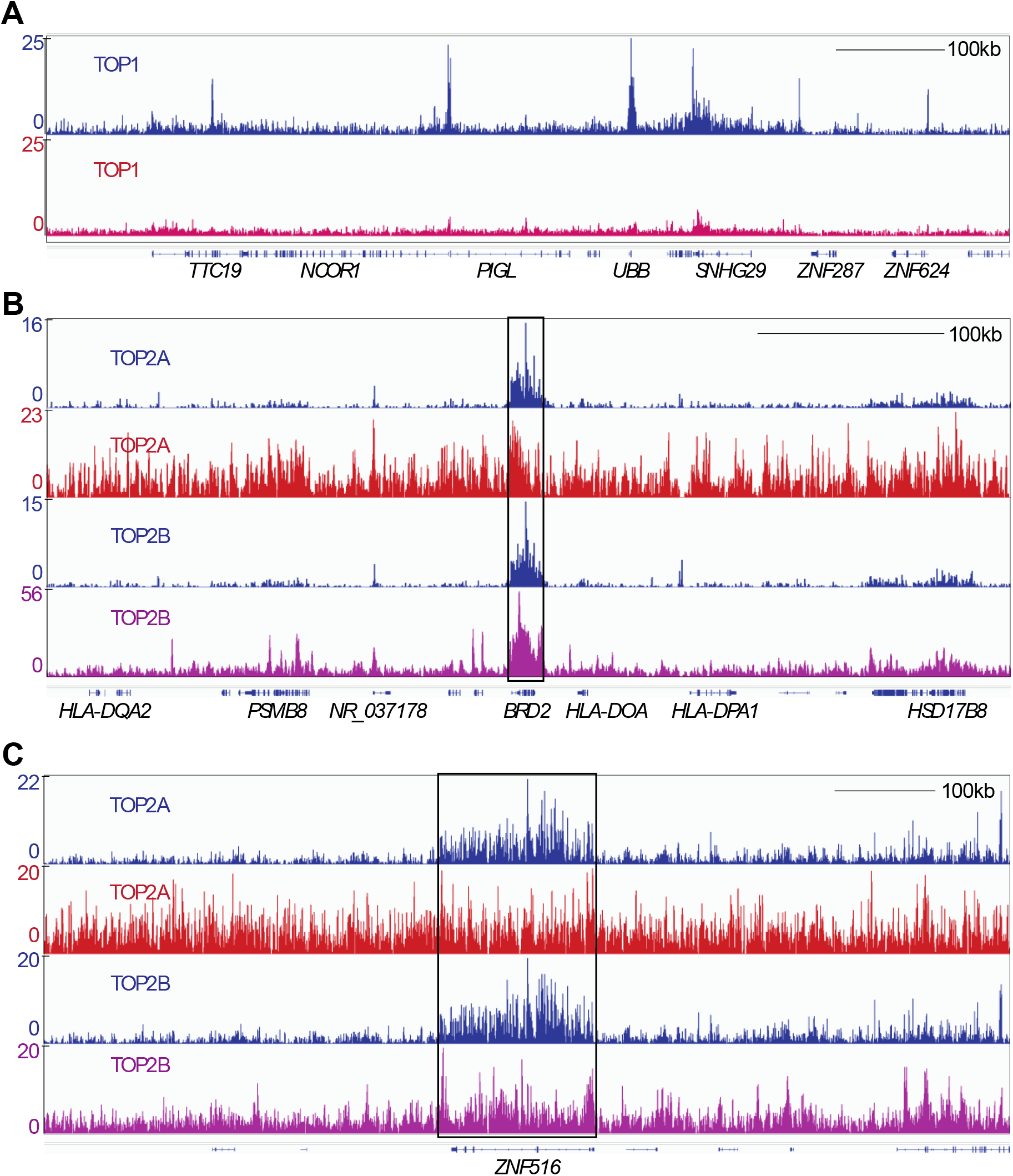
Comparison of our CUT&Tag sequencing methods with traditional ChIP methods. CUT&Tag signals are shown in blue and those from published prior methods in red. Representative example of IGV (Integrative Genomics Viewer) tracings of TOP1, TOP2A and TOP2B. Each topoisomerase signal was auto scaled with the scale shown on the left. A, Comparison of TOP1 binding signals. B, Comparison of TOP2A and TOP2B binding signals. The BRD2 gene regions is boxed. C, Comparison of TOP2A and TOP2B binding signals. The ZNF516 gene region is boxed.

**Fig. 3.**
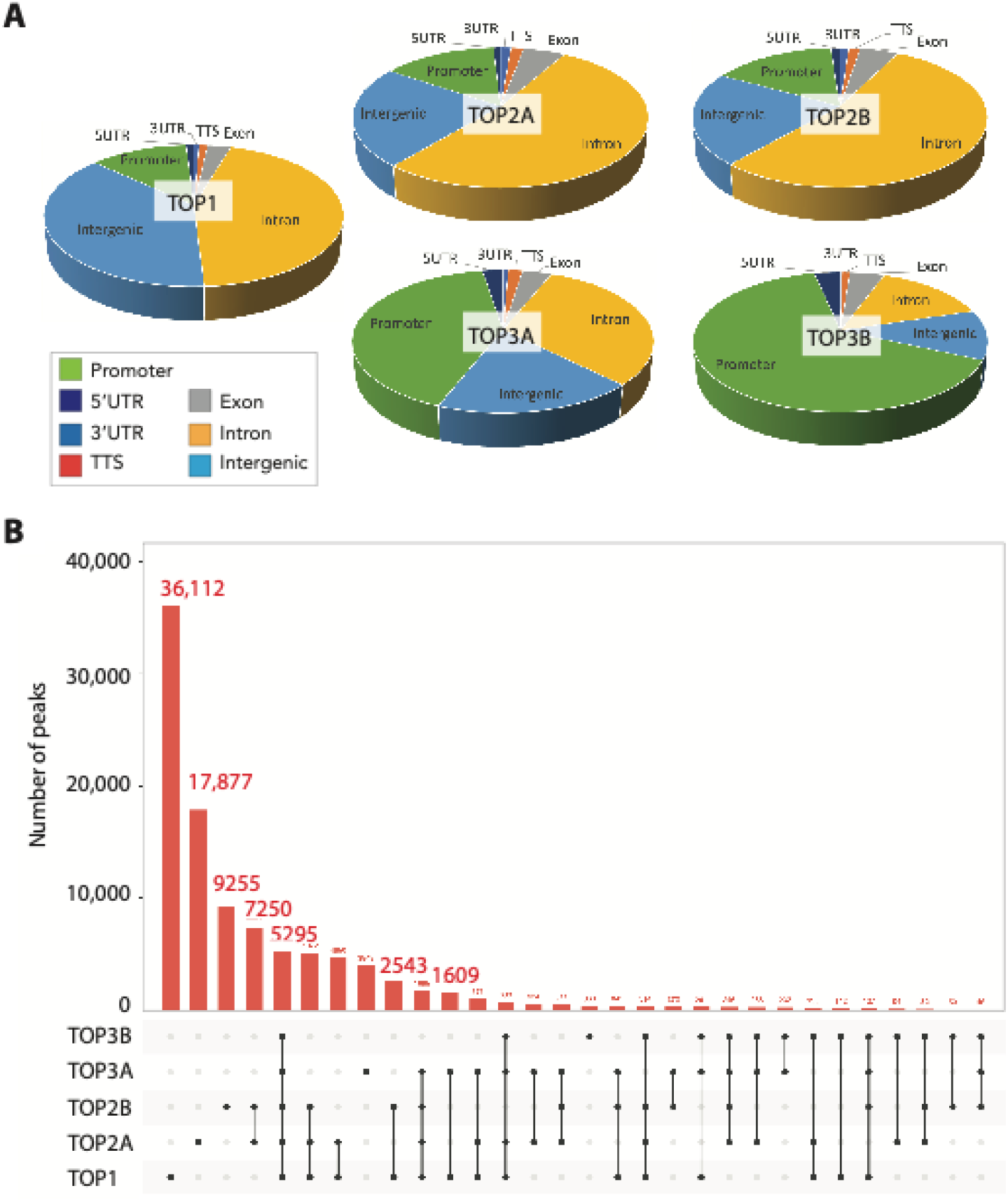
Topoisomerase binding signal analysis. A, Pie graph of topoisomerase binding signal distribution for all nuclear DNA topoisomerases (TOP1, TOP2A, TOP2B, TOP3A, and TOP3B). Abbreviations: 5UTR, 5’ untranslated region; 3UTR, 3’untranslated region; TTS, transcription termination region (site). B. Common and unique genomic peaks of all nuclear topoisomerases.

### Comparison of CUT&Tag with the prior ChIP-seq methods

The locations of topoisomerases on chromosomes have been studied previously^11^ and for TOP1, TOP2A, and TOP2B ChIP-seq data are available for each enzyme mapped individually^12–15^. We used publicly available data with the best signal to noise ratio and aligned the signals with our topoisomerase binding sites obtained using our CUT&Tag protocol.

For TOP1, we used GEO accession GSM2058666 (cell line: HCT116, Top1_ChIP-seq). In Figure 2A, our CUT&Tag TOP1 signals are shown in blue in comparison with the signals obtained from GSM2058666 in red in a representative region of chromosomes 17p11. Although both methods could map TOP1 binding to the same genomic locations (SNHG29, UBB and NCOR1 genes), the CUT&Tag method displayed improved signal to noise ratio (Figure 2A). For TOP2A and TOP2B, we used GEO accession GSM4213396 (cell line: hTERT RPE-1, TOP2A-CHIP-seq) and GSM4205700 (Cell line: MCF7, TOP2B-CHIP-seq), respectively. In figure 2B-C, we aligned representative TOP2A and TOP2B signals from our CUT&Tag method (in blue) with the TOP2A signals from GSM4213396 (in red) and TOP2B signals from GSM4205700 (in purple) in chromosome 6p21 (panel B). The representative IGV tracings for BRD2 (figure 2B) and ZNF516 (figure 2C) genes on chromosomes 6p21 and 18q23 (panel C) show that our CUT & TAG method produces the clearest signals both for TOP2A and TOP2B with better signal to noise ratio (SNR). Overall, these observations establish CUT&Tag as a reliable and sensitive method for mapping the genomic binding sites of topoisomerases.

### Overall genomic distribution of the nuclear topoisomerase binding sites

By compiling the overall topoisomerase binding signals, we were able to compare the signal distribution for the five nuclear DNA topoisomerases (TOP1, TOP2A, TOP2B, TOP3A, and TOP3B) over the different functional regions of the genome (Figure 3A). TOP1 peaks were the most abundant (Figure 3B) and were mainly in introns and in intergenic regions. The TOP2A and TOP2B peaks were similarly distributed and prominently localized in introns. They were equally distributed in promoters and intergenic regions. TOP3A and especially TOP3B showed high enrichment in promoter regions.

We also determined all the common and unique genomic peaks of across all nuclear topoisomerases (Figure 3B). For the unique peaks, TOP1 had the highest number of peaks (36,112) while TOP3B had the least (360). The common peaks for all 5 topoisomerases were 5295 and tended to be in promoter regions. TOP2A and TOP2B had the most overlapping peaks with 7250 such peaks. By contrast, TOP3A and TOP3B had only 232 peaks in common with TOP3A showing more than 10 times more peaks than TOP3B, which produced the smallest number of detectable peaks (360 peaks; Figure 3B) primarily localized in promoter regions (see Fig. 3A and Fig. 4A).

**Fig. 4.**
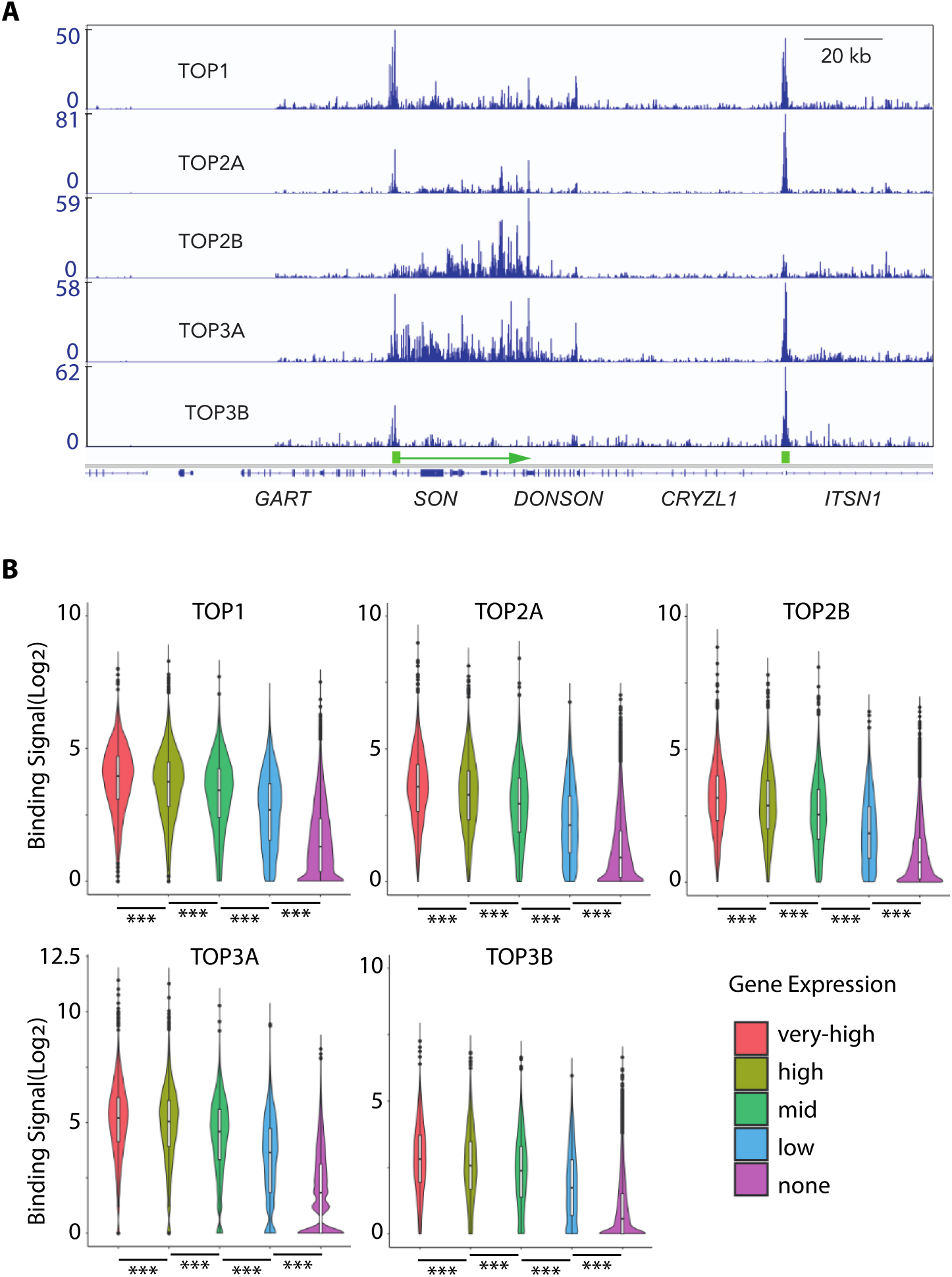
Topoisomerase binding signals in relationship with transcription. A, Representative example of IGV (Integrative Genomics Viewer) image of the distinct binding pattern of all five nuclear DNA topoisomerases. Each topoisomerase signal was auto scaled with the scaled indicated on the left. B, Topoisomerase binding signals positively correlate with gene expression. Genes are grouped based on their transcription levels: very-high, high, mid, low and no expression. All showed highly significant differences. With paired t test, the p-value was < 2.0E-16, and with Chi-square test, the p-value was <0.001.

### Distinct binding patterns of the individual topoisomerases

Comparative analyses of the IVG signal tracings for each of the topoisomerases showed distinct binding pattern for each topoisomerase within individual gene regions. Figure 4A shows a representative genomic DNA binding pattern in a segment of chromosome 21q22. For TOP1 and TOP2A, the CUT&Tag peaks were prominently at the promoter with additional smaller peaks in the body of the highly transcribed *SON* gene. By contrast, TOP3A and TOP2B binding was observed mostly in the gene body, and the TOP2B signals were more intense and shifted to the 3’-end of the gene (Figure 4A). For TOP3B, consistent with the global gene distribution analysis (see Fig. 3A), binding signals were primarily confined to the promoters of *SON* and *ITSN1*. For TOP3A the signals were clearly identified both in the promoter and gene body regions (Figure 4A). However, as discussed below the gene body signals of TOP3A for long genes are confined to the first 20 kb region (see the *DYRK1A* gene example in Figure 6).

### Topoisomerase binding is correlated with gene expression and highly transcribed regions

Because topoisomerases are known to be associated with chromatin remodeling, a dynamic process during transcription (see review^17^), we next asked whether overall topoisomerase binding intensity per gene is related to the expression of each gene. To do so we crossed the published gene expression data of HEK293^46^ (SRR710092.2) (the cells used for our analysis) with our topoisomerase genomic mapping data. After stratification of all the genes into five categories: very-high, high, mid, low, and not-expressed genes, we observed that the very highly expressed genes have the most topoisomerase binding signals, followed by high, mid and low expression genes, whereas the non-expressed gene have the least topoisomerase binding signals (Figure 4B). We computed the statistical significance value using paired t-test and Chi-square test. With paired t-test, the p-value was < 2.0E-16. Similarly, with the Chi-square test, the p-value was <0.001, indicating a highly significant positive correlation between overall gene binding signals for all the five nuclear topoisomerases and the expression level of individual genes (Fig. 4B).

Further analyses revealed that highly transcribing genes coding for transcription-associated proteins constituting nuclear speckles such as the splicing factor *SON*^47–50^ showed prominent topoisomerase signals (Fig. 4A). We noticed that only some actively transcribed genes possessed topoisomerase signals, which was reminiscent of that specific actively transcribed regions can interact with nuclear speckles^47,48^. To investigate the potential relationship between transcription nuclear speckles (i.e. transcription splicing hubs) we compared the mapping of SON, a core component of nuclear speckles^49,50^ with the mapping of topoisomerases. Figure 5A shows a representative tracing of SON determined by TSA-Seq^49^ and topoisomerase binding across the P arm of chromosome 2. Notably, both SON and the topoisomerases showed overlapping patterns, although our CUT&Tag signals for topoisomerases were from HEK293 cells and the SON TSA signal was from K562 cells (GSE81553_SON)^49^. The wide SON peaks matched clusters of topoisomerase peaks, which themselves were similar to each other (Fig. 5A). This point is emphasized in the overlay of the different tracings (bottom of Fig. 5A). Together, these result shows that topoisomerase binding matches transcription/splicing hubs.

**Fig. 5.**
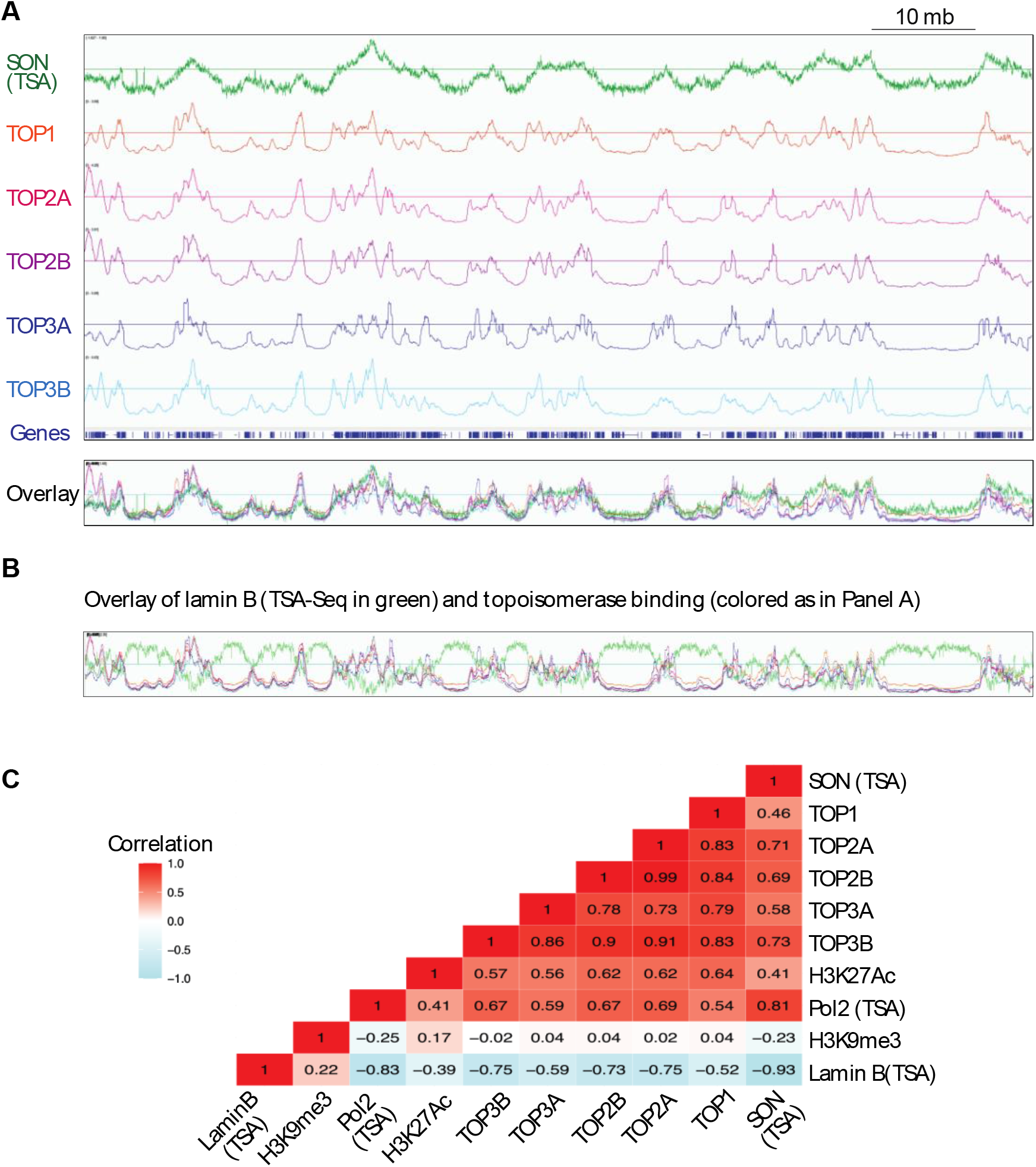
Topoisomerase binding signals coincide and are positively correlated with active of transcription. A, Tracing of the P-arm of chromosome 12 (1-90 mb) showing the binding of SON (a core nuclear speckle component, RNA processing center) (TSA) as indicator of nuclear speckle distribution across the whole chromosome arm. The binding signals from all nuclear topoisomerases coincides with SON binding. B, The binding signals from all of nuclear topoisomerases are inversely correlated with the tracing of Lamin B (TSA), a marker of transcriptional repression in the same chromosomal area as in panel A. C, Overall correlation of the overall peak distribution of the nuclear topoisomerases and markers of active and inactive transcription H3K27Ac, Pol2(TSA), H3K9me3 and Lamin B (TSA).

To confirm this result, we compared the topoisomerase binding clusters with the binding sites of Lamin B (nuclear lamina component), which are transcription-suppressed chromatin sites. Figure 5B shows a representative tracing overlay for the p-arm of chromosome 2. Contrary to SON, the Lamin B contour pattern was opposite to the topoisomerase clusters (Figure 5B). This result shows that topoisomerase signals are negatively correlated with transcriptional repression signals, and altogether this data suggest that topoisomerases preferentially bind to highly transcribed chromatin regions.

To further match our topoisomerase CUT&Tag signals with positive and negative markers of active transcription, we chose H3K27Ac, SON, and Pol2 as markers for active transcription and H3K9me3 and Lamin B as markers for repressed transcription. For H3K27Ac (GSM2711409)^51^ and H3K9me3 (GSM5330293)^52^, the data were from ChIP-seq, and for SON (GSE81553_SON), Pol2 (GSE81553_Pol2) and Lamin B (GSE81553_LaminB), they were from TSA-seq (estimating cytological distances of chromosome loci genome-wide relative to a particular nuclear compartment)^49^. After converting the files to BigWig format, we used the bigWigCorrelate tool from the UCSC Genome Browser utilities to compute the Pearson correlation coefficients between the signal values in the input BigWig files. Figure 5C shows high positive correlation for the H3K27Ac, SON and Pol2 signals (all in red) with the topoisomerase binding signals and with each other, while the Lamin B signals were negatively correlated with the topoisomerase/H3K27Ac and SON signals. H3K9me3 (heterochromatin marker) did not show any significant correlation with the topoisomerase binding signals (Figure 5C). Additionally, the highest correlation was observed between TOP2A and TOP2B (Pearson correlation coefficient: 0.99) consistent with the common binding features of these two topoisomerases (Fig. 5C). This result is consistent with the analyses shown in Figure 3.

### Arresting DNA replication suppresses TOP3A binding

To examine whether progressing DNA replication affects the genomic binding sites of topoisomerases, we treated cells with aphidicolin, an established DNA polymerase inhibitor^53^ before performing CUT&Tag (Fig. 6A). Figure 6B shows representative tracings for chromosome 21q22 demonstrating that aphidicolin reduces the signals of TOP2A, TOP2B and TOP3A (here shown for the *DYRK1A* gene) with the largest suppressing effect on TOP3A (Fig. 6B, bottom two tracings). Simultaneously, aphidicolin enhanced the TOP3A, TO2B and TOP2A signal in the promoter regions (marked with short green box and arrow to the left of *DYRK1A*) (Fig. 6B). Notably, aphidicolin had minimal effect on TOP1 and TOP3B signals (Fig. 6C).

**Fig. 6.**
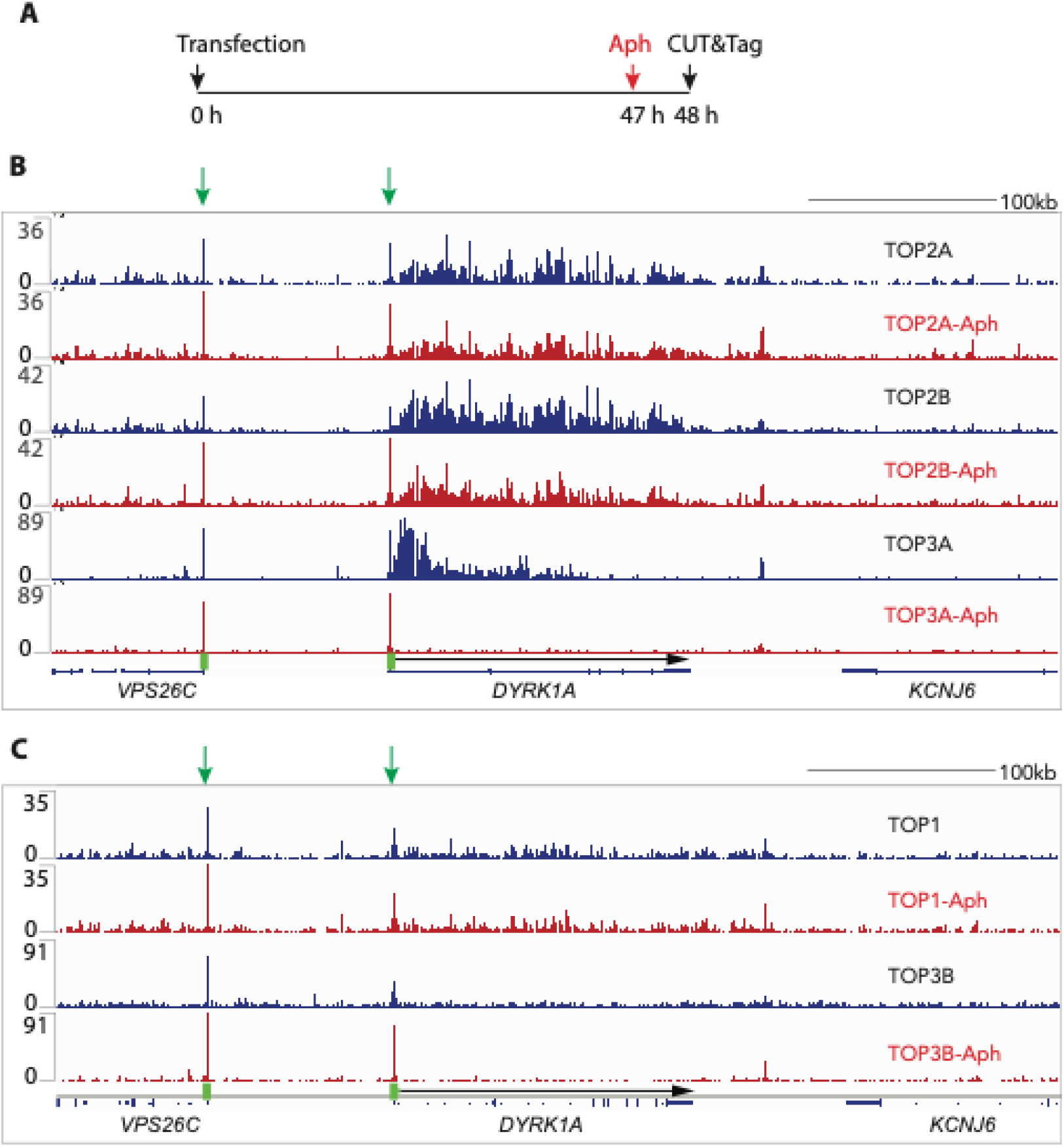
Replication-dependency of topoisomerase binding. A, Outline of the experiment. B, Representative example of IGV (Integrative Genomics Viewer) image of all nuclear topoisomerase binding signals. Tracings from untreated cells are in blue and from aphidicolin-treated (1 µM for 1 hour) in red. Each pair is plotted with same scale (labeled on the left) for comparison. The DYRK1A promoter region is labeled as green vertical rectangle. The gene body is indicated as a black arrow.

Because TOP3A was among the 5 topoisomerases, the most affected by aphidicolin, and we had previously shown by RADAR assays the global suppression of TOP3A binding by aphidicolin,^35^ the last part of our study focuses on the genomic distribution of TOP3A binding upon replication arrest by aphidicolin.

First, we analyzed in which genes the TOP3A binding was most affected by aphidicolin. To do so, we compared the TOP3A binding signals associated with each gene in aphidicolin-treated vs. untreated cells (Fig. 7A). While most genes showed suppression of TOP3A signal, only a limited number of genes showed some increase, such as the *TEDC1* gene (Fig. 7A and Fig. S2A). The genes where TOP3A signals are most suppressed by aphidicolin (Fig. 7A) are RNA binding/regulatory genes, as demonstrated by Gene Set Enrichment Analysis (GSEA) using the Enrichr software (https://maayanlab.cloud/Enrichr/enrich#, Go biological process) for the genes whose TOP3A signals were at least 100 cpm (count per million) and showed at least 3-folds decrease in aphidicolin-treated cells. The 691 genes (highlighted area in Figure 7A) that passed these criteria, among which RNA processing related genes occupied the top 6 categories. (Fig. 7B).

**Fig. 7.**
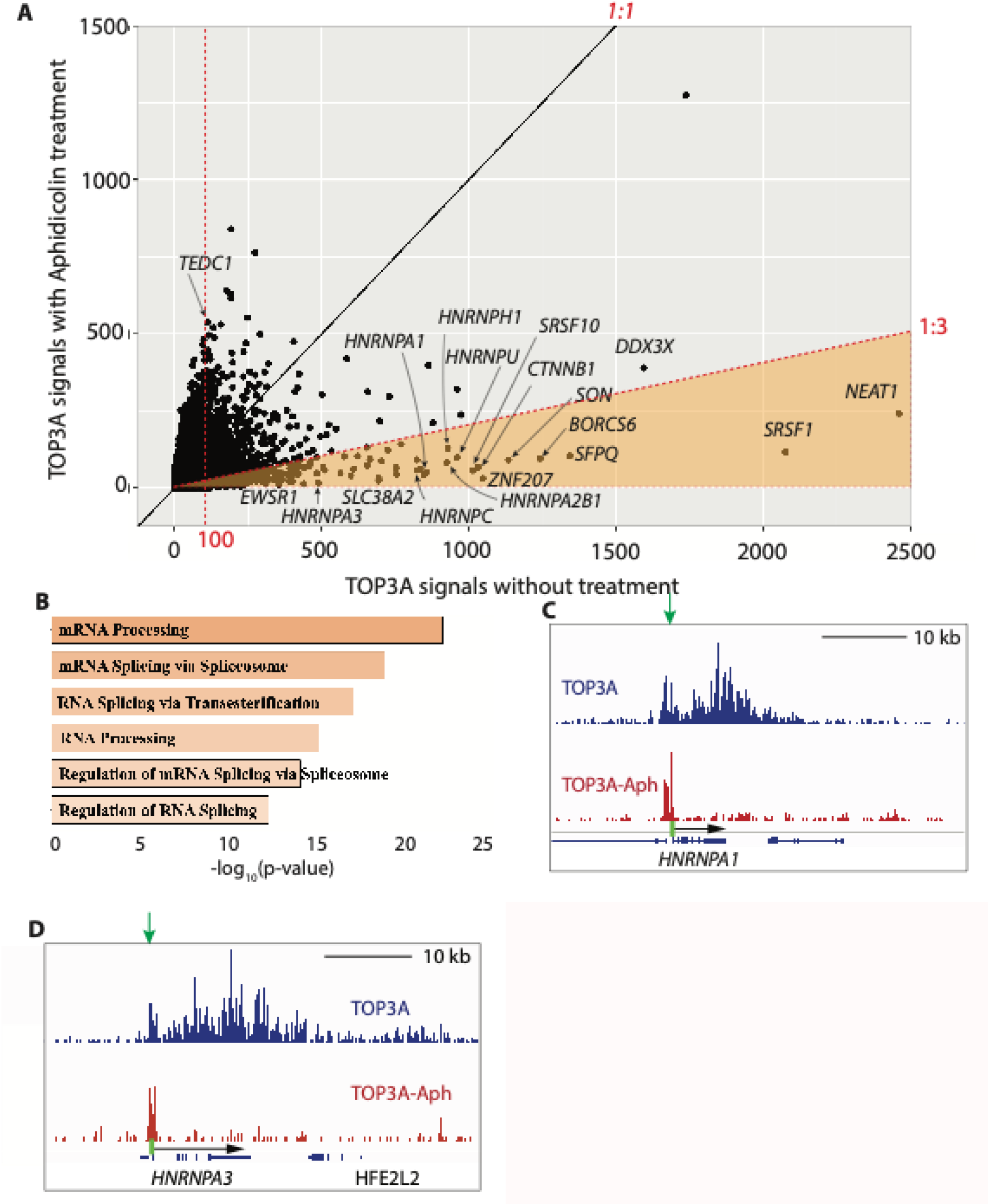
Replication-dependent TOP3A signals. A, Overall TOP3A binding signal plot with emphasis on genes with suppressed signals by aphidicolin. The area of TOP3A signaIs over 100 (rpm) and signal ratio of untreated over aphidicolin-treated over 3 is highlighted in orange. It includes 691 genes. **B,** GO biological process generated with Enrichr for the 691 genes whose TOP3A signal was suppressed by aphidicolin. C and D, Representative examples of IGV image of genes with replication-dependent TOP3A signals in gene body. Promoter regions are labeled as green rectangles. Gene bodies are indicated as arrows.

Among the genes where TOP3A binding was most reduced by aphidicolin, genes associated with nuclear speckles (such as *SRSF1*, *SON*, *HNRNPA1*, *HNRNPA*) showed the most striking effect of aphidicolin (Fig. 7A). Representative tracings showing suppression of TOP3A signals in gene bodies upon aphidicolin treatment for the *HNRNPA1* and *HNRNPA3* are shown in Figures 7C and 7D and for *SRSF1* gene in Supplementary Figure 2B. Together, these results reveal that TOP3A binding tends to be preferentially associated with highly transcribed genes coding for RNA binding/regulator proteins and that this TOP3A binding is rapidly reversed by arresting replication by aphidicolin, consistent with our recent results using RADAR assays^35^.

### Preferential binding of TOP3A in the 5’-region of transcribed genes: the 20 kb rule

Since TRCs can be both HO and CD, depending on the relative directions of replication and transcription, we studied the direction of potential TRCs. OK-seq replication-forward-direction (rfd) was used to determine the direction of DNA replication^39^. Since DNA replication may start from different sites in different cell lines, the replication directions (as determined by OK-seq) that were common in four cell lines (BL79, IMR90, K562, and TF1)^39^ were used as consensus. We manually listed the first 101 genes with prominent TRCs. Among these 101 genes, there were 20 different genes with potential HO-TRCs, 35 genes with CD-TRCs, and the rest of genes contained regions of both HO- and CD-TRCs for different cell lines. Both directions of TRCs caused topological stress and accumulation of topoisomerase signals (Fig. S3). The situation is clearly different from bacteria where the CD-TRCs are less detrimental^54^, resulting in minor topological problems. Thus, our results suggest that the CD-TRCs in human cells poses similar threat as the HO-TRCs in terms of replication-dependent recruitment of TOP3A.

Next, we analyzed the distribution of the topoisomerase signals within the genes showing potential TRCs as defined as replication-dependent TOP3A signals (see Fig. 7 and above). Since TOP1 and TOP3B signals did not respond to replication blockage, we also examined in parallel TOP2A, TOP2B and TOP3A signals as potential indicators of TRCs and topological stress. For small genes such as *SRSF3* (Fig. 8A, top panel), the topoisomerase signals were distributed across the whole gene body and even extended beyond the 3’-end. For longer genes, we found that the TOP2A and TOP2B signals covered the whole gene length while the TOP3A signals covered only about 20 kb from the 5’-end of the genes (see example of the *MED13* gene, Fig. 8, bottom panel). We refer to this phenomenon as the “20 kb rule of TOP3A binding in CD-TRCs”. Additional examples for the 20 kb rule are given in Figure S4. These results suggest that the long genes with CD-TRCs require TOP2A and TOP2B for resolution of topological stress throughout their gene bodies, whereas TOP3A is selectively acting in the first 20 kb of gene bodies to resolve TRCs.

**Fig. 8.**
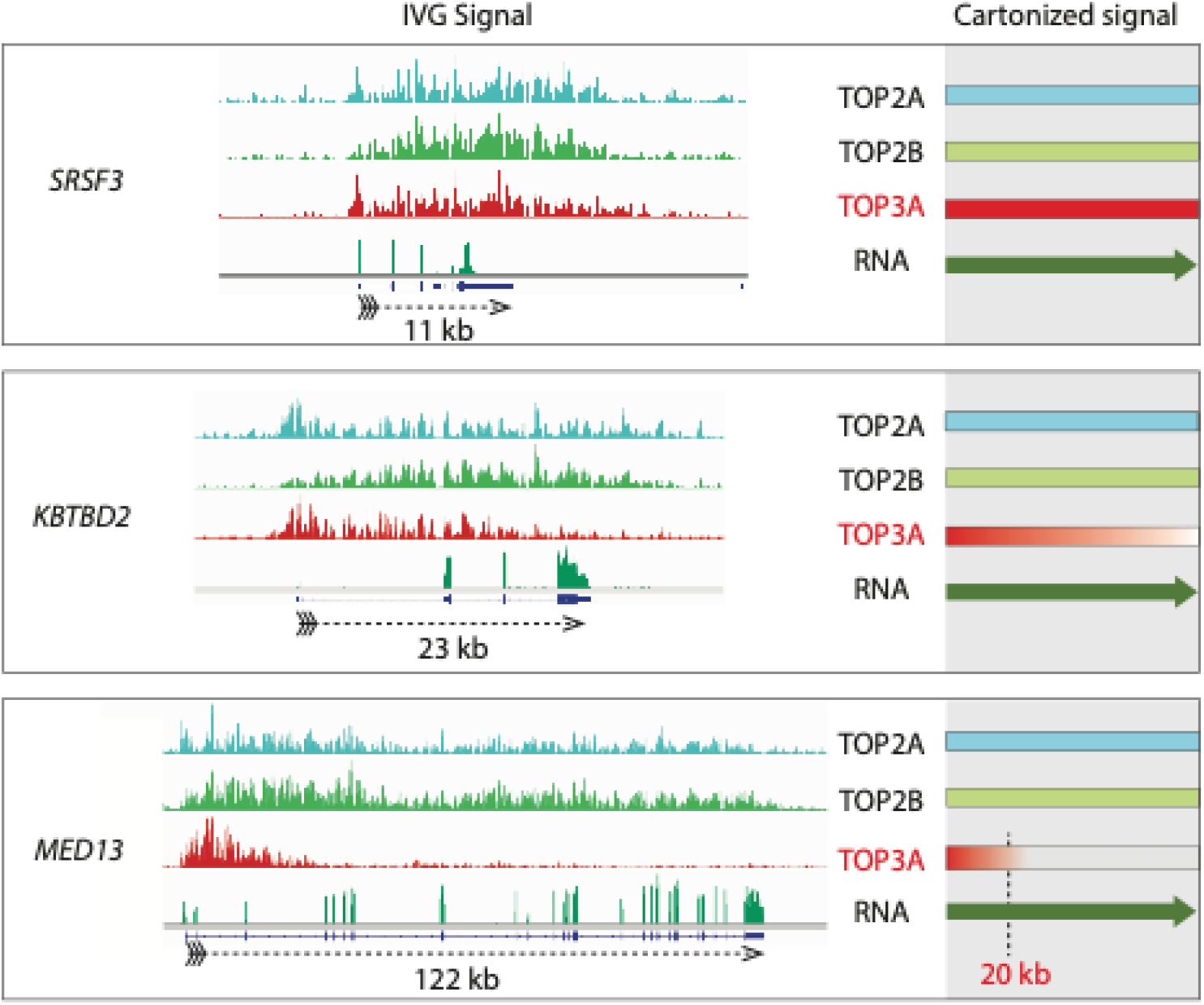
The TOP3A binding 20-kb rule. Representative IGV-(Integrative Genomics Viewer) tracings for three genes of increasing length (from top to bottom). TOP2A and TOP2B signals cover the full length of the transcribed genes whlie TOP3A signals cover about 20 kb at the 5’-end of the transcribed genes. The right panels are schemes to highlight the 20-kb rule.

## DISCUSSION

Traditional genome-wide binding signals of TOP1^12^, TOP2A ^13^, and TOP2B^14,15^ by ChIP-seq have relatively low signal to noise ratio (SNR) requiring the analyses to focus on specific regions of the genome with high signal often boosted by treatment with topoisomerase poisons. To improve the SNR, we applied a new CUT&Tag method (see Fig. 1) in human cells transfected with self-trapping constructs of DNA topoisomerases^22,35,44,45^. Utilizing this single NGS technique, we mapped the genomic locations of all five different nuclear topoisomerases with improved SNR. For TOP1, TOP2A and TOP2B, binding signals are consistent with previous studies (see Fig. 2). Notably, our study is the first to map the TOP3A and TOP3B binding sites across the whole genome.

Comparison of the genome-wide distribution of the topoisomerases shows similarities and distinctive features (see Fig. 3). We find that TOP1 displays the largest number of binding peaks, and that the binding peak frequency was by decreasing order: TOP1>>TOP2A>>TOP2B>>TOP3A>>TOP3B. Not unexpectedly peak distribution was most similar for TOP2A and TOP3B (see Fig. 3A). TOP1 peaks were characteristically high in non-coding regions (introns and intergenic regions) while TOP3A and even more TOP3B peaks were prominent in promoters (see representative tracings in Figures 2 and 4-7 and Supplemental Figures S1-4).

A common characteristic of the topoisomerase binding sites was their association with active transcription (see Fig. 4B) with preferential concentration in highly transcribing nuclear speckle regions (see Fig. 5). The correlation between the topoisomerase signals and nuclear speckles suggests the spatial arrangement of topoisomerases in nuclei. Indeed, we find that genes encoding nuclear speckles (*SRSF1*, *SON*, *HNRNPA1*, *HNRNPA3*, etc.) are most prominent sites of topoisomerase binding (see Fig. 7A). The link between the nuclear speckle genes and their products is consistent with two observations. First, some gene products can regulate their own transcripts. For instance, SRSF1 (SF2/ASF) and SRSF2 (SC35) negatively autoregulate their expression to maintain homeostatic levels through alternative splicing^55,56^. Second, nuclear speckle components bind to the DNA of other nuclear speckle genes. While exploring our ChIP-seq dataset for the speckle-components, we found that HNRNPH1, HNRNPK, HNRNPL and other speckle gene products could bind to the beginning of the *HNRNPA3* gene (not shown), which is one of the most affected speckle-component-genes affected by replication inhibition (see Fig. 7D). High transcription rate within a small confined nuclear compartment may cause more TRCs, resulting in more topological constraints that require TOP3A. As Speckle-associated domains remain largely conserved^57^ among different cell lines, we used the SON signals (TSA-seq) published from K562 cells, and found high correlation between the SON signals and the topoisomerase signals from HEK293 cell as well. To complement this picture, we observed that the transcriptional repression marker Lamin B is negatively correlated with the topoisomerase binding signals. These observations lead us to hypothesize that nuclear speckles (and other phase-separated nuclear domains) may set up an overall chromosomal environment encompassing coordinated DNA structures, RNAs and proteins (beyond transcription factors and enzymes) activities. Among them, our data suggest that topoisomerases are important components to monitor and coordinate the DNA and/or RNA structures for their normal functions.

Our study is the first to map the genome-wide distribution of both human type-IA topoisomerases TOP3A and TOP3B. As noted above, we find that the TOP3B peaks are strikingly less frequent that those of TOP3A and the other topoisomerases (see Fig. 3). We believe this is likely related to the fact that TOP3B is a dual DNA and RNA topoisomerase with prominent localization and activity in the cytoplasm and polysomes^58^ while its nuclear activities are to enforce genome stability by limiting R-loops^59^.

By contrast, we observed that TOP3A signals have the unique characteristics to be prominent at the promoters of highly transcribed genes and that among the long genes, they cluster within 20 kb of their 5’-regions (see Figs. 8 and S4). Most notably, these TOP3A signals are replication-dependent and suppressed by arresting replication with aphidicolin (see Figs. 6, 7 and S2). This result further validates our recent observations measuring TOP3A-DPCs by RADAR assay^35^. Additionally, we find that replication-dependent TOP3A signals increase with the number of intron/exon junction (Fig. S5). Together, these results imply that TOP3A plays a previously unrecognized role in being recruited to regions of transcription-replication conflicts (TRCs) both in head-on (HO-TRCs) and co-transcriptional (CD-TRCs) collisions (see Fig. S3). Thus, we propose the model represented in Figure 9 linking TOP3A to TRCs. How and why TOP3A is recruited to TRCs remains to be elucidated at the molecular level and is beyond the scope of the current study. One possibility is that TRCs lead to the rotation of replication forks as they encounter transcription complexes and that such rotation generates precatenanes behind the revolving replication forks with regions of single-stranded DNA where TOP3A can act as a decatenase^17^.

**Fig. 9.**
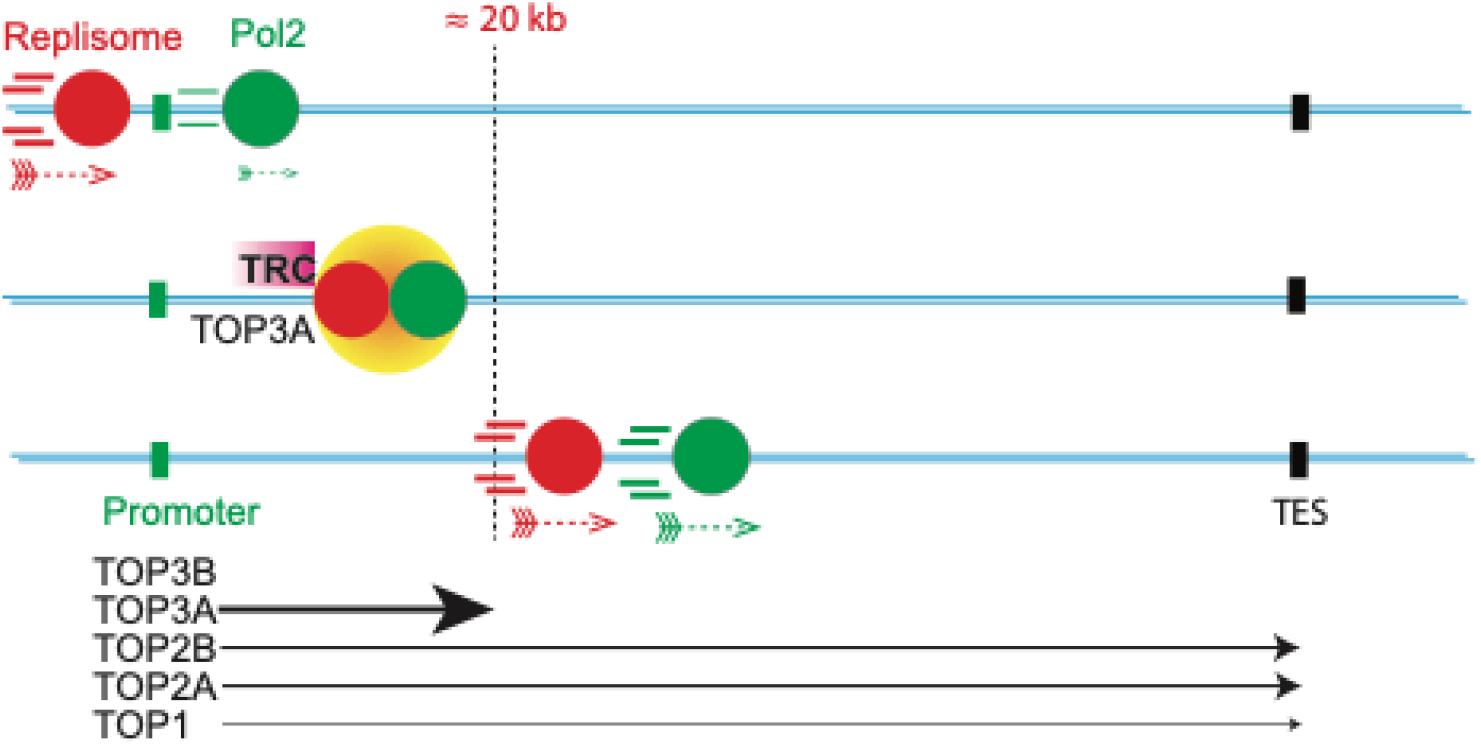
Proposed role of topoisomerases in Transcription Replication Conflicts (TRCs). Outside of transcribing genes, replisomes move at higher speed than RNA polymerase (Pol2), In highly transcribed genes, replisomes catch up with Pol2 complexes and TRCs recruit topoisomerases to alleviate topological problem (see Discussion). TOP3A is recruited possibly to resolve precatenanes arising from replisome rotation. If the genes are long, after traveling 20 kb of distance, fully active Pol2 moves at the similar speed as the replisome. TOP2A and TOP2 remove the excessive supercoils and no extra TOP3A is needed.

## DATA AVAILABILITY

The CUT&Tag data have been deposited in the GEO database under accession number GSE269841

## SUPPLEMENTARY DATA

Supplementary Figures

## Authors’ Contributions

H.Z., Y.S., S.S., and L.K.S. designed and conducted the experiments, H.Z., L.S.P., and A.D. performed the informatics analysis, Y.P. supervised the study. H.Z., Y.S., S.S., L.K.S., and Y.P. wrote the manuscript.

## Acknowledgments

Our studies are supported by the Center for Cancer Research, the Intramural Program of the NCI (Z01-BC006150, to Y. Pommier).

