## Supplemental Figures 1-5 for "Genome-wide Mapping of Topoisomerase Binding Sites Suggests Topoisomerase 3α (TOP3A) as a Reader of Transcription-Replication Conflicts (TRC)"

### Supplementary Figures

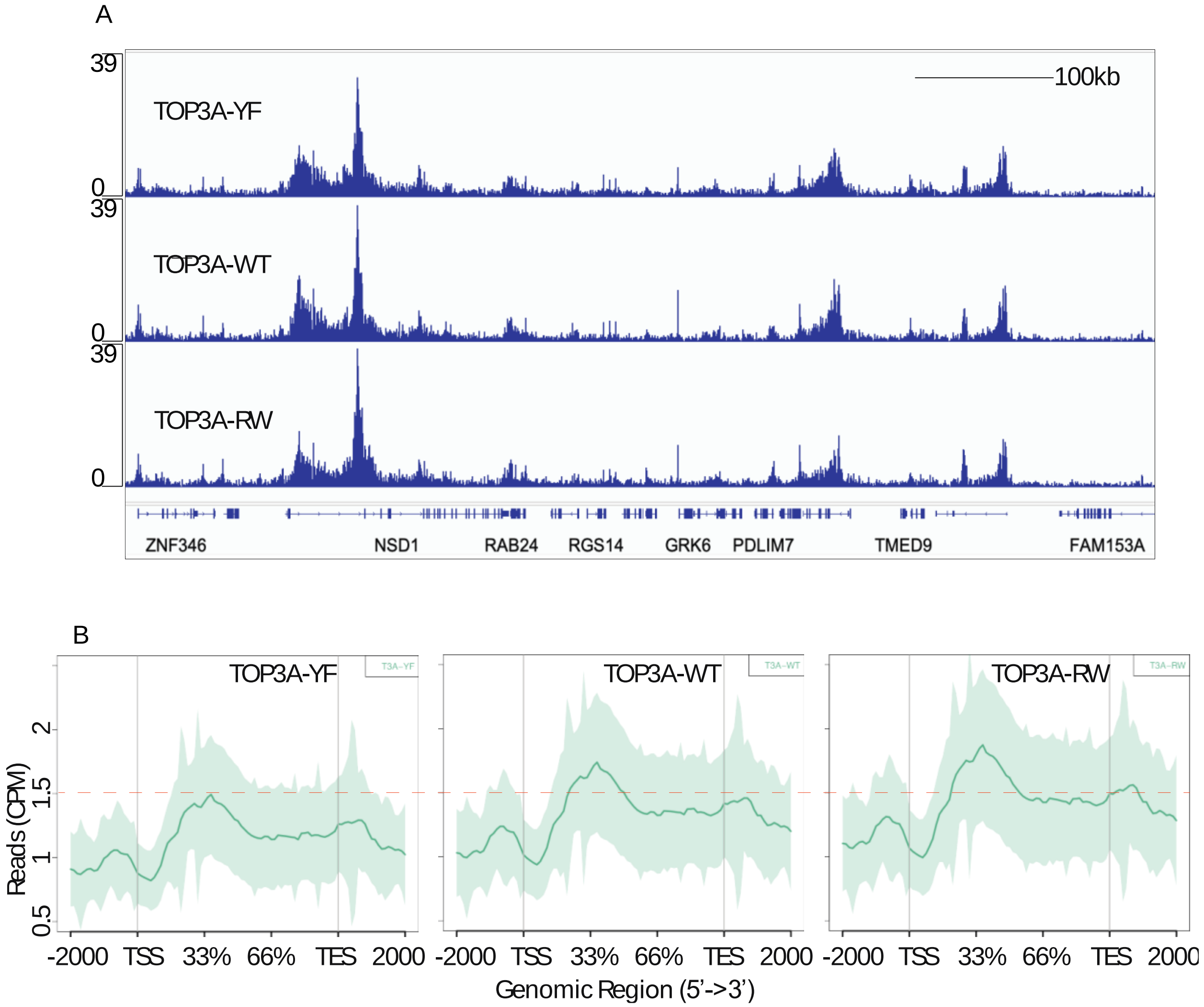


Fig. S1. Binding signals of the 3 forms of TOP3A: WT: wild-type, YF: catalytic dead by mutation of the catalytic tyrosine Y362 to phenylalanine (F), and RW: mutation of arginine R364 to tryptophane (W) induces the self-trapping of TOP3A on DNA. **A**, Representative example of IGV (Integrative Genomics Viewer) image of the binding pattern of Y362F, WT and R364W. **B**, Overall binding patterns of Y362F, WT, and R364W.

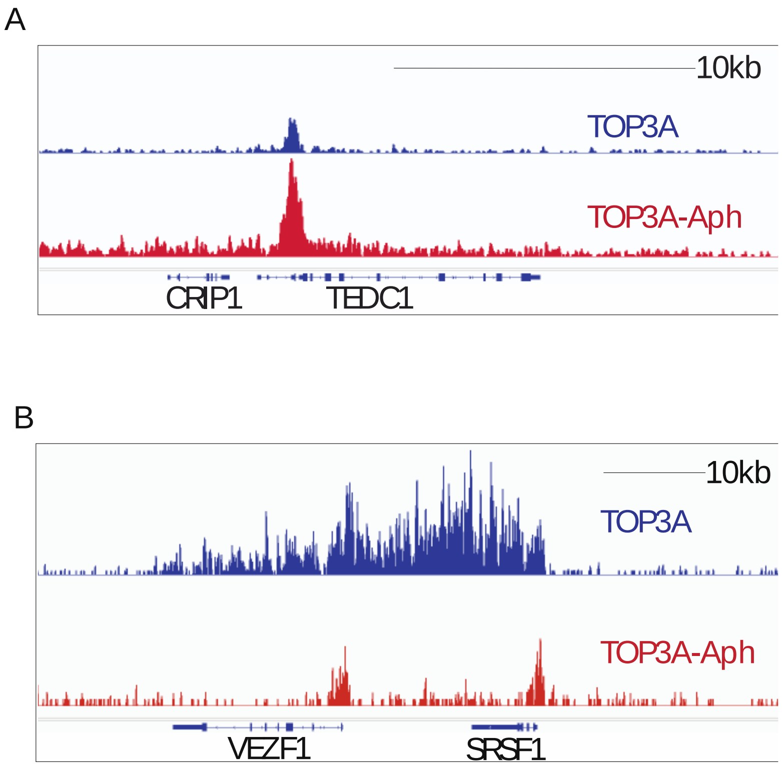


Fig. S2. Additional examples of genes with replication-dependent TOP3A signals. **A**, As indicated in Figure 7A, *TEDC1* is an example of the small number of genes whose TOP3A signals are increased by aphidicolin. **B**, *SRSF1* is an example of genes with suppression of TOP3A signals by aphidicolin (see Fig. 7A).

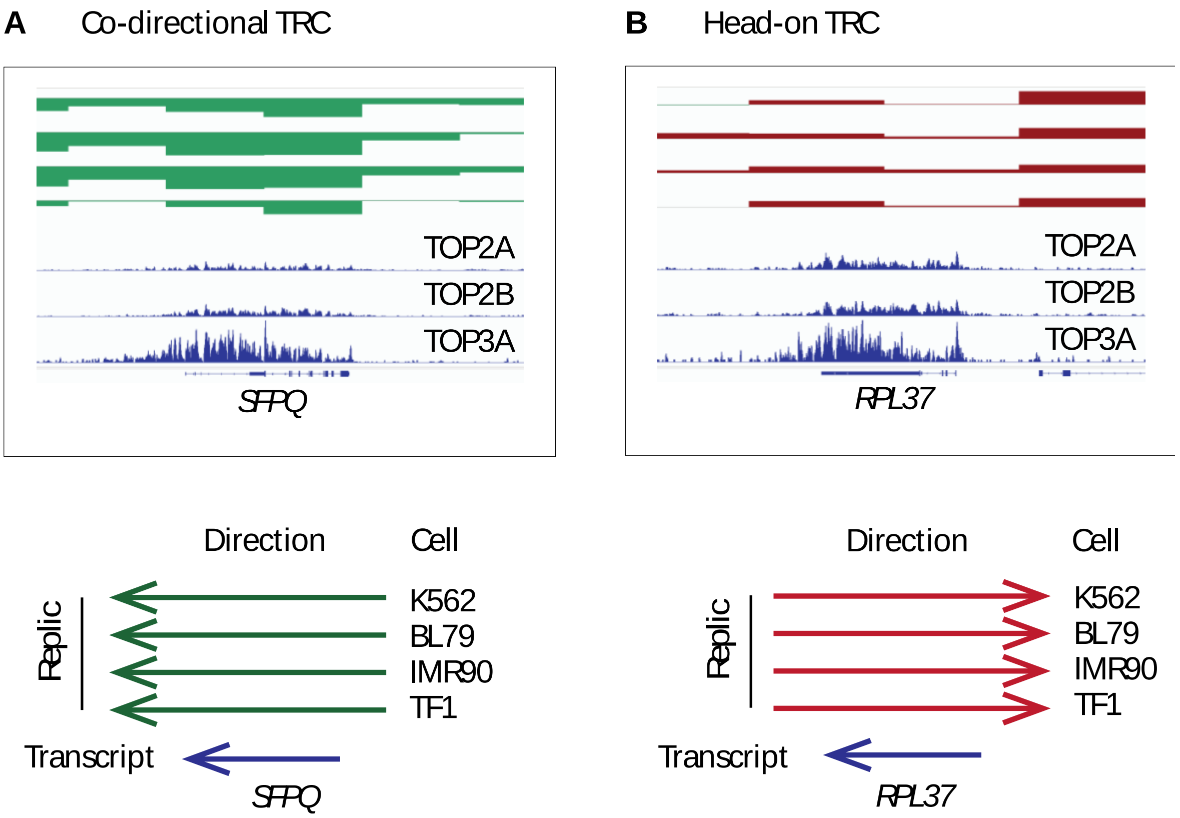


Fig. S3. Topoisomerase binding signals occurs on genes with both CD-TRCs and HO-TRCs. **A**, *SFPQ* is an example of co-directional TRC with recruitment of TOP2A, TOP2B and TOP3A. Replication and transcription move in the same directions based on published data in 4 human cell lines. Both replication and transcription are shown moving from right to left with the OK-seq labeled in green. **B**, *RPL37* is an example of HO-TRC that recruits TOP2A, TOP2B and TOP3A Replication and transcription are from opposite directions based on published data in 4 human cell lines, as indicated.

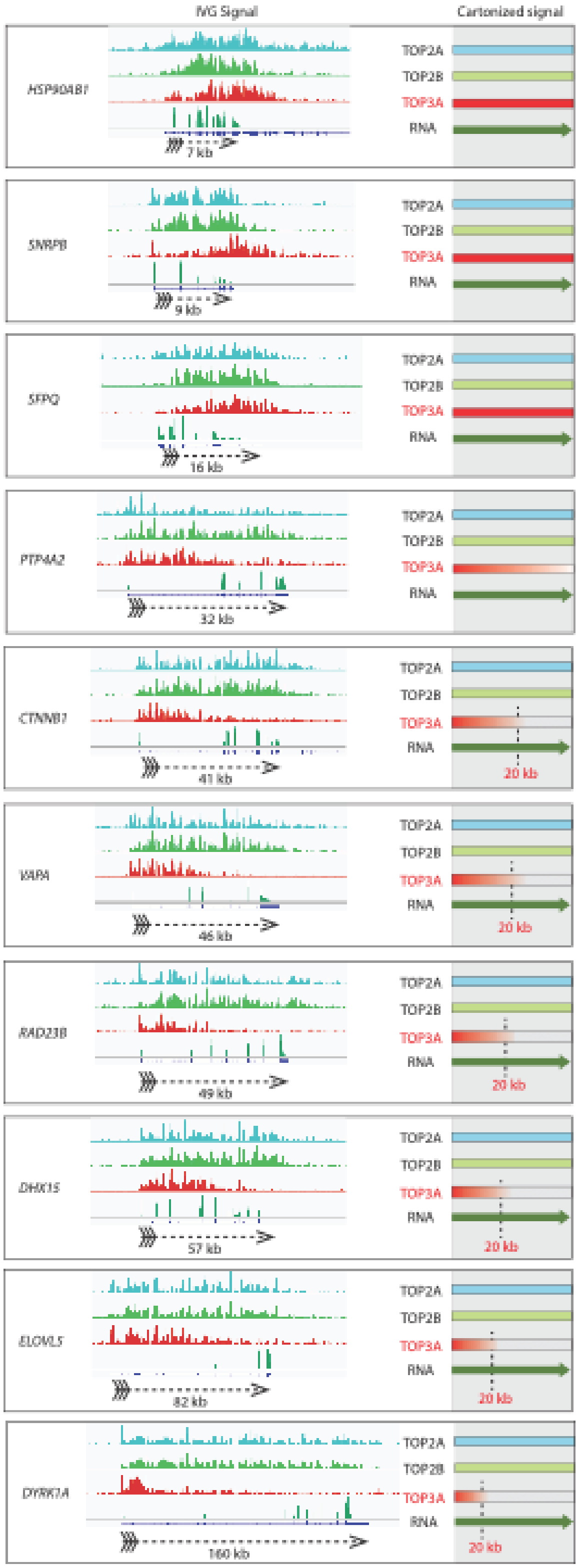


Fig. S4. Representative examples of the 20-kb rule complementing the data shown in Figure 8. Gene are ordered by increasing length from top to bottom with lengths indicated under dashed arrows. Right panels shaded in gray are schematic representations.

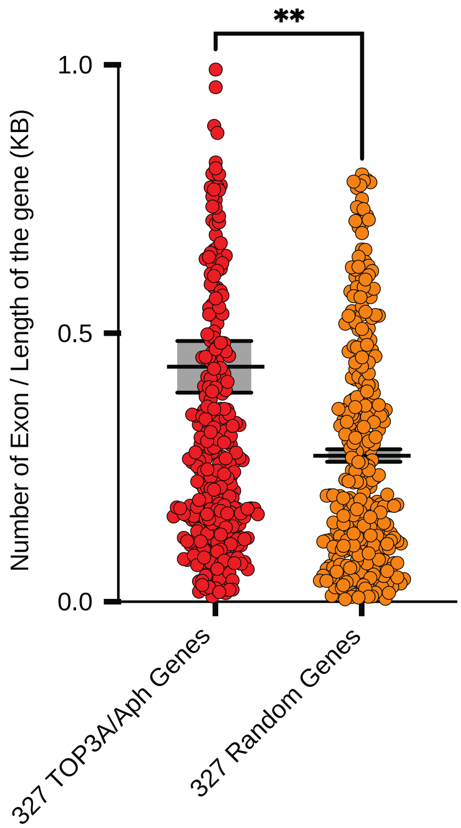


Fig. S5. Higher exon/intron junction density for genes with TOP3A signals in comparison with random selected genes. Exon frequency (numbers of exons / length of the gene in kb) of 327 genes whose TOP3A signals were suppressed by replication inhibition compared with a set of 327 genes chosen randomly. The t-test shows a significant increase in replication-dependent TOP3A signals as a function of intron/exon juntions.
